# Freshwater soundscapes: a cacophony of undescribed biological sounds now threatened by anthropogenic noise

**DOI:** 10.1101/740183

**Authors:** Rodney A. Rountree, Francis Juanes, Marta Bolgan

**Affiliations:** The Fish Listener, East Falmouth, Massachusetts, United States of America; Biology Department, University of Victoria, Victoria, British Columbia, Canada; Laboratoire de Morphologie Fonctionnelle et Evolutive, Institut de Chimie, Université de Liège, Liège, Belgium

**Author notes:** Corresponding author (RR).

## Abstract

The soundscape composition of freshwater habitats is poorly understood. Our goal was to document the occurrence of biological sounds in a large variety of freshwater habitats over a large geographic area. The underwater soundscape was sampled in freshwater habitat categorized as brook/creek, pond/lake, or river, from five major river systems in North America (Connecticut, Kennebec, Merrimack, Presumpscot, and Saco) over a five-week period in the spring of 2008. Over 7,000 sounds were measured from 2,750 minutes of recording in 173 locations, and classified into major anthropophony (airplane, boat, traffic, train and other noise) and biophony (fish air movement, also known as air passage, other fish, insect-like, bird, and other biological) sound categories. Anthropogenic noise dominated the soundscape of all habitats averaging 15 % of time per recording compared to less than 2 % for biological sounds. Anthropophony occurred in 79 % of recordings and was mainly due to traffic and boat sounds, which exhibited significant differences among habitats and between non-tidal and tidal river regions. Most biophonic sounds were from unidentified insect-like, air movement fish, and other fish sound sources that occurred in 57 % of recordings. Mean frequencies of anthropogenic noises overlapped strongly with the biophony, and comparisons of spectra suggest that insect-like and air movement sounds may be more susceptible to masking than other fish sounds. There was a significant decline in biodiversity and biophony with increasing ambient sound levels. Our poor understanding of the biophony of freshwater ecosystems, together with an apparent high temporal exposure to anthropogenic noise across all habitats, suggest a critical need for studies aimed at identification of biophonic sound sources and assessment of potential threats from anthropogenic noises.

## Introduction

Fish sounds were first seriously studied in North America by Abbot [1] who lamented that “the little fishes of our inland brooks and more pretentious denizens of our rivers are looked upon as voiceless creatures…” and concluded that “certain sounds made by these fishes are really vocal efforts….”. However, Stober [2] was the first to describe the underwater noise spectra in relation to freshwater fish sounds. He also pointed out the critical lack of data on the ambient noise and fish sounds in freshwater systems. Fortunately, after long neglect there has been a recent surge in interest in the potential impacts of anthropogenic noise on freshwater ecosystems [3–7], but efforts to understand such impacts are hampered by the paucity of data on the natural soundscape composition. The paucity of data on the sound production of freshwater fishes was highlighted in our recent review of the literature which found that sounds have been reported in only 87 species in North America and Europe, but detailed descriptions of sound characteristics are known for only 30 species [8].

To our knowledge previous studies of the biological soundscape of freshwater habitats in the New England region of North America include species-specific studies [9–11] and our survey of the Hudson River [12]. Martin and Popper [6] conducted a survey of the noise levels in the vicinity of the Tappan Zee Bridge in the Hudson River, but did not examine biological sounds. In a pilot study of the fish sounds in the Hudson River we discovered a high diversity of known and unknown biological sounds [12]. At a location in New York City, arguably one of the world’s most impacted locations, we also found a high diversity of mostly unknown biological sounds at night after boat noise declined [12]. The value of reporting information on unknown biological sound occurrence was emphasized when we later discovered that one of the unknown sounds in the Hudson River was produced by the invasive freshwater drum [9]. We followed up the Hudson River study with this pilot study in 2008 intended to document the soundscape of a wide range of habitats within the New England region. Unfortunately, at that time we knew too little about the characteristics of freshwater fish sounds to accurately process the recordings. We, therefore, subsequently conducted a series of surveys to identify some of the most common fish sounds and document the importance of Fast Repetitive Ticks (FRTs) and other air movement sounds made by fishes in a companion paper [10]. Based on this new understanding of freshwater fish sounds, we were able to reprocess recordings from the 2008 study in order to classify sounds into broad biological sound types. The primary goals of this study were to document the occurrence of biological sounds in a large variety of freshwater habitats over a large geographic area and to determine the relative contribution of broad categories of biological and anthropogenic sounds to the underwater soundscape composition.

## Materials and Methods

### Study area

The underwater soundscape of freshwater habitats was sampled in a roving survey along five major river systems in the New England region of North America over a five-week period from 30 April to 29 May 2008 (Fig 1). Rivers surveyed included the 653 km Connecticut River (30 April to 3 May, N = 32), 188 km Merrimack River (12 – 16 May, N = 43), 270 km Kennebec River (17 – 23 May, N = 53), 219 km Saco River (26 – 28 May, N = 31) and 42 km Presumpscot River (28 – 29 May, N = 14). Sound recordings were made from shore within the main stems of each river from their origin in the mountains or major lakes to their outlets to the sea, except for the Connecticut River where only the lower 200 km were surveyed. In addition, sound recordings were made from other habitats such as major tributaries, small sluggish creeks and rivers, woodland streams, and ponds and lakes in each river region (S1 Fig). Access to the water for sampling was often difficult and usually required sampling within 500 m of a bridge (45 %) or dam (12 %), and sometimes involved scrambling down steep ravines, over rocks, or short hikes through woodlands. All locations were photo-documented. Although both lotic and lentic habitats sampled represent a continuum, for comparison purposes, brooks and streams were combined with small sluggish rivers or creeks into a “brook/creek” habitat category, small ponds and large lakes were combined into a “pond/lake” category, and tributary and main stem-river sites were combined into a “river” category (S1 Fig).

**Fig 1.**
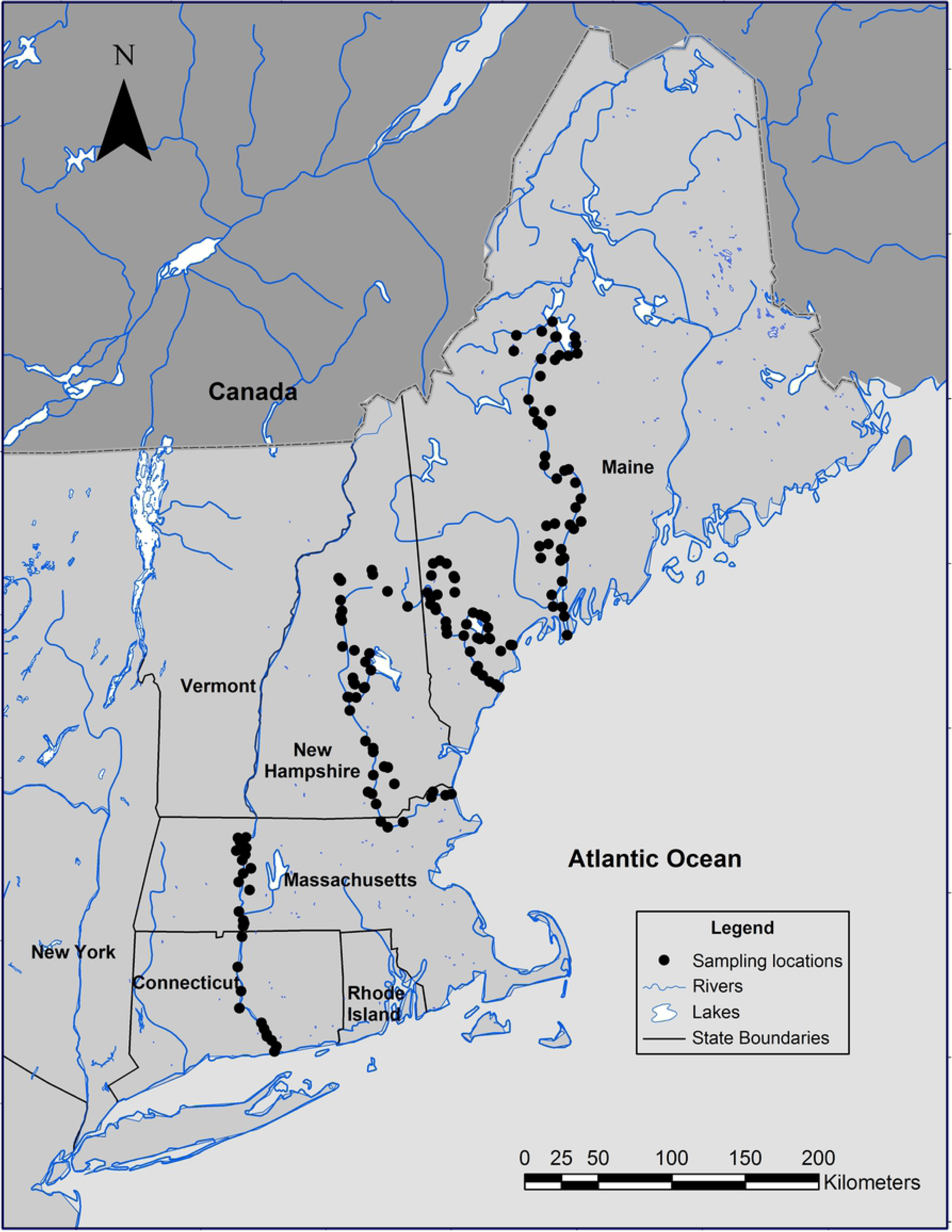
Study area. Locations where sounds were recorded between 30 April and 30 May 2008.

A total of 173 sites were successfully sampled during the survey (S1 Table). Additional sites were attempted but could not be sampled due to poor environmental conditions or equipment failure. Sampling was avoided during poor weather (high winds and/or rain). Sampling occurred during the day between the hours of 0930 and 1900 (N = 148), and at night between 1900 and 2300 h (N = 25). Night sampling sometimes encompassed daylight hours prior to sunset. Recordings were made from 1 to 49 minutes duration (mean = 12.5) during day and 3 to 119 minutes (mean = 35.5 minutes) during night sampling (S1 Table). Night recordings were longer on average than day recordings because we expected increased biological activity around sunset based on previous experience [7, 9–10, 12], and to avoid travel in unfamiliar and often remote locations after dark.

### Acoustic data

Acoustic data were captured from a low noise, broadband (flat response from 16 - 44,000 Hz), cylindrical hydrophone with 30 m of cable (model C54XRS, Cetacean Research Technology, Seattle, Washington) at 44.1 kHz (16 bit) with a MOTU Ultralite, bus powered firewire audio interface to a laptop computer using SpectraPro332 Professional Sound Analysis software (Sound Technology, Inc.). Calibrated hydrophone frequency response data were provided by the manufacturer between 20 Hz and 14 kHz. Additional total system calibration data were provided for the audio interface at zero and half-full gain settings. Sound from either a dubbing microphone or a second hydrophone was captured to a second channel in the recordings. When used the second hydrophone was an uncalibrated, variable (dial) gain Aquarian model (Aquarian Audio Products, Anacortes, WA, USA). A scrolling spectrogram and waveform were monitored continuously while listening to the recording in real-time in the field. Written and oral notes on sounds together with potential environmental, anthropogenic and biological sound sources were also recorded throughout each recording. Additional post-processing of acoustic signals was conducted by listening to all recordings in their entirety while simultaneously viewing the sound’s spectrogram (1024 FFT, Hanning window, 50% overlap) and waveform with Raven Pro 1.5 acoustic software (Bioacoustics Research Program 2014).

Most anthropogenic sounds (anthropophony) were positively identified in the field, including airplane, boat, traffic (bridge crossing or road), train and other noise. Boat sounds were divided into sounds of running boats, boats at idle, and other boat sounds (engine cranking, pumps, etc.). Fishing sounds were those made by recreational fly and spin-cast fishers and included lines hitting the water surface and sinking, lure retrieval, weights dragging on the bottom, etc. Other noise included miscellaneous construction sounds (e.g., hammer tapping) lawn mowers, fish/depth finders, etc.

Biological sounds were largely unidentified and subjectively classified into broad categories based on our previous experience [7, 9–10, 12]. Sound type categories included: Fast Repetitive Ticks and other air movement sounds [sensu 10], other fish, insect-like, surface, bird and other biological sounds. Bird sounds were recorded underwater from sounds arising above water. Other fish sounds included more conventionally recognized fish sounds such as drumming and stridulation sounds. Because we have previously observed that the sounds made by fishes when they jump or gulp air at the surface can be useful identification markers, and may potentially have inter- or intra-biological functions [10], we include them in a “Surface” sound category. Other biological sounds included beaver (tail slaps) and seal sounds, as well as unknown snapping sounds (snap sounds made by Atlantic sturgeon, *Acipenser oxyrinchus*, were included in the other fish category). Unknown sounds deemed to be biological but which could not confidently be placed within one of the biological categories were also placed in the other biological sound category. The unclassified sound category included bubble-like and gurgle sounds because a type of air movement sound labeled “gurgle” described for salmonids, cannot yet be reliable distinguished from natural sounds of gas release from the sediment [10]. Additional sound category groupings were used in some analyses, including air movement (FRT + other air movement), fish (air movement + other fish), biophony (fish + insect-like + bird + surface + other biological), boat (running boat + idle boat + other boat), anthropophony (airplane + boat + fishing + traffic + train + other noise), and unclassified (gurgle + bubble-like + unknown).

Within each separate recording unique biological sound types were annotated and subjectively labeled (e.g., FRT-A, FRT-B) and the number of unique biological sound types counted (i.e., biological sound diversity per recording). However, while biological sound categories were consistent across all recordings, individual biological sound types were not, due to the high variability of sound characteristics, and diversity of habitats sampled, so total biological sound diversity could not be determined. For example, a sound labeled a “bark” and placed in the other fish category in one recording might have different acoustic characteristics from a “bark” in another recording, making it difficult to determine if the observed differences were due to variability or different sources. However, in that case the sound could still be confidently placed in the “other fish” category. In addition, two 5 s clips from each recording were annotated to represent ambient sounds, herein defined as the background sound when no individually recognizable biological or anthropogenic sounds were observed [sensu 15].

Total received sound pressure levels (RSPL) were calculated in SpectraPro332 for each ambient clip and each recording was arbitrarily assigned into one of four Root Mean Square (RMS) [13] sound level categories (90-95, 96-100, 101-105, and >105 dB RMS RSPL re 1 μPA). Sound levels were not estimated in some recordings due to mechanical noise on the calibrated hydrophone, strong flow, cable strumming, or other factors. Many locations (28 %) apparently exhibited electromagnetic field (EMF) levels high enough to introduce EMF noise into the recording and were excluded. Average spectra of each sound category, including ambient, were calculated from a subsample (S2 Table) of representative clips (Hanning, FFT 4096, 50% overlap, frequency resolution 11.7, PSD normalized). In addition, we compared the received sound level of selected biological sound categories to ambient sound levels by calculating the received sound level above the ambient in each recording and then averaging across all subsamples for that category. For example, after linearizing, the spectra of the ambient sound were subtracted from each FRT subsample from each of the ten locations where the FRT subsamples were selected, and then all the resulting spectra were averaged over all 23 FRT subsamples.

Acoustic measurements of all sound types were made in Raven of selected parameters including duration, peak frequency and frequency bandwidth [14]. The “percent frequency of occurrence” (not to be confused with acoustic frequency parameters) of sounds was determined as the number of recordings containing a sound, divided by the total number of recordings. Sound rate was calculated as the number of sounds in each recording for each category standardized to number per minute. Biological diversity is reported as the number of sound types per recording. Sound “percent time” was calculated for each sound category by summing the durations of all sounds in the category and dividing by the recording duration and is expressed as a percent of the recording time per record. Temporal overlap among biological and anthropogenic sounds was determined by counting the number of sounds that overlapped completely, or partially, in time (i.e., that occurred at the same time in the recording).

### Data analyses

Many environmental factors that potentially influence soundscape composition such as habitat category, diel period, river position and river system were statistically confounded, but were compared based on various subsamples of the data in an exploratory examination of their potential to influence the soundscape composition. The exploratory nature of these comparisons is emphasized given the “snap-shot” nature of the data collection and lack of control of environmental variables such as time-of-day, temperature, season (although all data were collected over a five-week period), amount of human development (ranging from remote wilderness to heavily populated urban areas), turbidity, water depth, and propagation distance. Because strong diel differences were expected, and night sampling was limited, all soundscape measurements (percent frequency, sound rate, and percent time) are reported separately for day and night recordings. Factors examined for possible influence on the soundscape composition included diel period, habitat type, position along the river gradient (examined separately for each river), river region (non-tidal and tidal reaches pooled over all rivers for day samples) and RMS category. Simple Chi square tests of the percent frequency of occurrence of sound types among data groups were used to suggest diel, habitat type and RMS sound level category effects. A one-way analysis of variance was used to test for potential single-factor effects (e.g., habitat category) after transforming to normalize.

## Results

A total of 4,825 biological, 1623 anthropogenic noise, and 834 unclassified sounds (Table 1) were measured from 2,750 minutes of recording in 173 locations (S1 Table). Examples of some of the most common air movement sounds are described elsewhere [10]. Examples of traffic, train, running boat and other boat noises are provided in S2-S5 Figs together with their corresponding sound files (S1-S4 Audio). An example of other fish, other air movement, and FRT sounds relative to traffic noise can be viewed in S2 Fig and heard in S1 Audio.

**Table 1.** Sound duration and peak frequency.

| Sound category | Duration (s) |  |  |  |  | Frequency (Hz) |  |  |  |  |  |
| --- | --- | --- | --- | --- | --- | --- | --- | --- | --- | --- | --- |
|  | Samples | Mean | SE | Min | Max | Samples | Mean peak | SE peak | Max peak | Mean IQR-BW | SE IQR-BW |
| **Biophony** |  |  |  |  |  |  |  |  |  |  |  |
| FRT | 135 | 3.03 | 0.34 | 0.16 | 31.53 | 125 | 1717 | 102 | 4969 | 586 | 46 |
| Other air movement | 1636 | 0.19 | 0.01 | 0.02 | 13.29 | 1506 | 1742 | 28 | 10359 | 607 | 14 |
| Other fish | 1807 | 0.32 | 0.03 | 0.01 | 32.90 | 1612 | 726 | 9 | 2625 | 303 | 5 |
| Insect-like | 631 | 2.21 | 0.10 | 0.02 | 29.62 | 238 | 2820 | 121 | 12188 | 776 | 94 |
| Bird | 57 | 2.29 | 0.39 | 0.11 | 16.02 | 54 | 2912 | 195 | 4781 | 752 | 79 |
| Surface | 197 | 0.84 | 0.04 | 0.06 | 4.60 | 188 | 906 | 35 | 3797 | 414 | 23 |
| Other biological | 362 | 0.20 | 0.04 | 0.01 | 13.05 | 325 | 1811 | 87 | 11438 | 933 | 56 |
| Subtotal | 4825 |  |  |  |  | 4048 |  |  |  |  |  |
| **Anthropophony** |  |  |  |  |  |  |  |  |  |  |  |
| Airplane | 11 | 27.58 | 3.66 | 14.76 | 51.18 | 9 | 307 | 130 | 984 | 120 | 24 |
| Fishing | 74 | 2.51 | 0.57 | 0.16 | 32.20 | 67 | 1483 | 230 | 8344 | 1133 | 244 |
| Running boat | 57 | 174.64 | 18.73 | 15.37 | 733.44 | 45 | 875 | 126 | 4266 | 1094 | 333 |
| Idle boat | 14 | 276.66 | 68.96 | 14.19 | 788.62 | 11 | 435 | 162 | 1406 | 464 | 138 |
| Other boat | 127 | 6.97 | 1.30 | 0.06 | 100.62 | 75 | 1159 | 90 | 3609 | 523 | 53 |
| Traffic | 1237 | 7.90 | 0.26 | 0.05 | 132.00 | 1070 | 225 | 19 | 4594 | 151 | 6 |
| Train | 22 | 44.53 | 29.81 | 0.06 | 654.99 | 5 | 459 | 103 | 563 | 84 | 50 |
| Other noise | 81 | 15.70 | 2.58 | 0.01 | 182.91 | 63 | 847 | 99 | 2906 | 373 | 48 |
| Subtotal | 1623 |  |  |  |  | 1345 |  |  |  |  |  |
| **Unclassified** |  |  |  |  |  |  |  |  |  |  |  |
| Gurgle | 313 | 1.24 | 0.11 | 0.09 | 18.13 | 296 | 1108 | 20 | 3797 | 229 | 10 |
| Bubble-like | 105 | 2.39 | 0.24 | 0.06 | 17.44 | 88 | 948 | 29 | 2109 | 222 | 21 |
| Unknown | 416 | 1.02 | 0.10 | 0.01 | 18.24 | 334 | 1616 | 69 | 10172 | 580 | 44 |
| Subtotal | 834 |  |  |  |  | 718 |  |  |  |  |  |
| **Ambient** |  |  |  |  |  | 159 | 297 | 34 | 1875 | 481 | 84 |
| **Total** | 7282 |  |  |  |  | 6270 |  |  |  |  |  |

Summary statistics for sounds pooled over all samples within sound categories. SE = Standard error of the mean, Min = minimum value, Max = maximum value, peak = frequency of greatest energy, IQR-BW = interquartile bandwidth.

In general, anthropogenic noises were 1 to 2 orders of magnitude longer in duration than biological sounds and exhibited consistent spectral overlap with them (Fig 2, Table 1, S6 Fig). FRTs, insect and bird sounds had the longest durations of the biophony averaging 2-3 s, while boat, plane and train sounds had the longest durations of the anthropophony averaging 28-277 s. Other fish sounds had the lowest peak frequency of the biophony, while insect and birds had the highest. Traffic sounds had the lowest mean peak frequency and fishing and other boat sounds had the highest peak frequency of the anthropophony. Train sound peak frequency was inflated by whistle sounds, and otherwise would have the lowest peak frequency (S6 Fig). Peak frequency of ambient sound was below that of most biological sounds but overlapped strongly with traffic and other anthropogenic noises (Fig 2, Table 1, S6 Fig).

**Fig 2.**
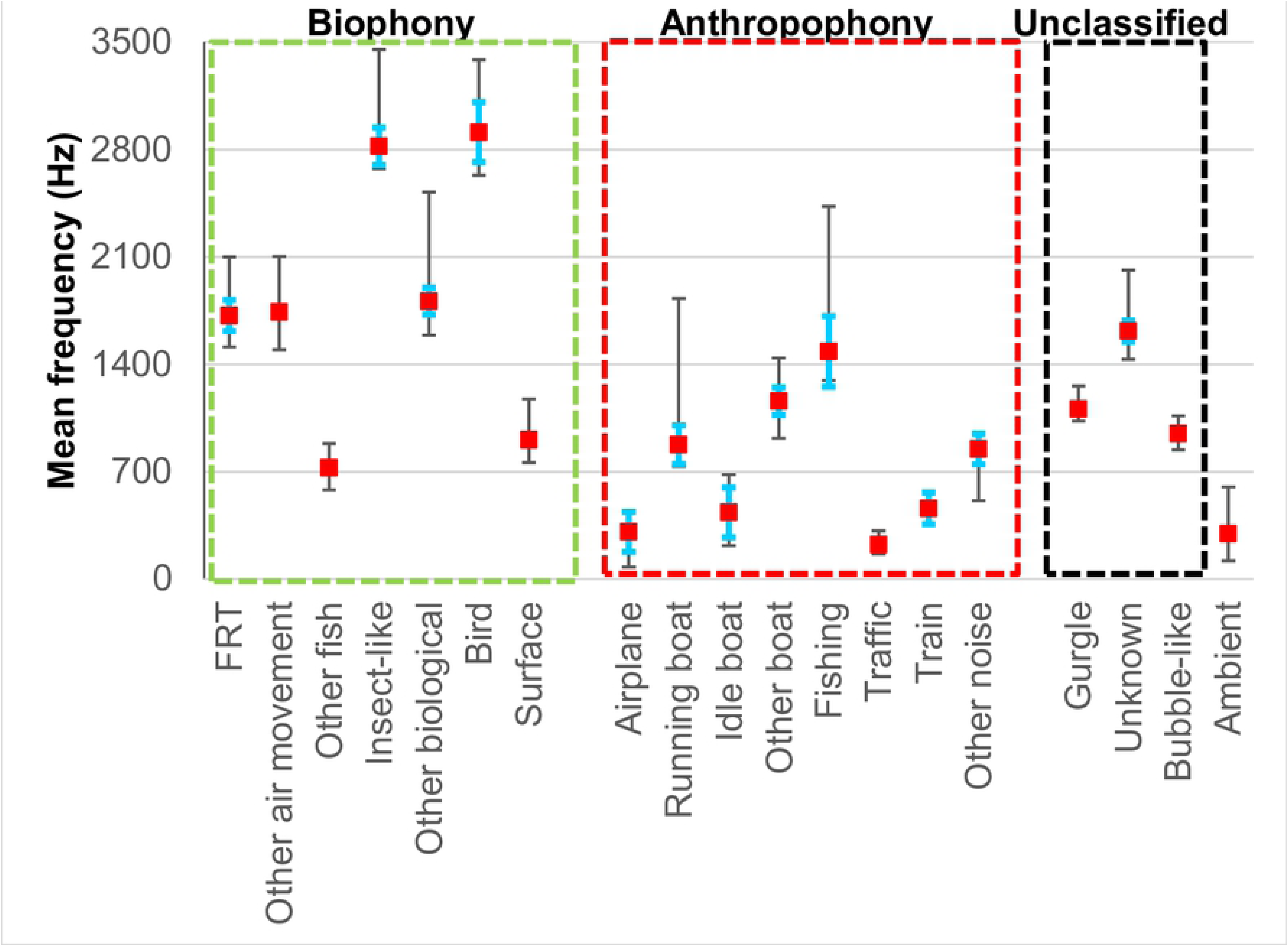
Sound duration and peak frequency. Comparison of acoustic characteristics among sound types and ambient sound. Square symbols mark the mean peak frequency, blue hats mark one standard error of the mean. The lower stem marks the mean first quartile frequency and the upper stem the mean 3rd quartile frequency (see Table 1).

Biophony occurred in 57 % of the recordings, while anthropophony occurred in 63 % (Table 2). Other air movement sounds were the most frequently occurring (39 %) component of the biophony followed by other fish (30 % of the recordings). Air movement sounds were dominated by high-frequency sounds similar to those previously described [10] for salmonids (46 %), alewife (*Alosa pseudoharengus*, Clupeidae) and alewife-like sounds (27 %), and white sucker-like (*Catastomus commersonii*, Catostomidae) sounds (8 %). Twenty-three percent of FRT sounds were attributed to alewife while the rest were unknown. Most of the other fish sounds were unknown, but 13% were catfish-like sounds most likely produced by brown bullhead (*Ameiurus nebulosus*, Ictaluridae).

**Table 2.** Comparison of sound occurrence between diel periods.

| Sound category | Day samples | Night samples | Total | Chisq P |
| --- | --- | --- | --- | --- |
| **Biophony** |  |  |  |  |
| FRT | 22% | 40% | 24% | * |
| Other air movement | 39% | 76% | 45% | *** |
| Air movement | 42% | 80% | 47% | *** |
| Other fish | 30% | 64% | 35% | *** |
| Fish | 48% | 84% | 53% | *** |
| Insect-like | 14% | 24% | 15% | ns |
| Other biological | 20% | 44% | 23% | *** |
| Surface | 14% | 56% | 20% | *** |
| Bird | 13% | 20% | 14% | ns |
| All biophony | 53% | 84% | 57% | *** |
| **Anthropophony** |  |  |  |  |
| Airplane | 4% | 8% | 5% | ns |
| Running boat | 13% | 24% | 14% | ns |
| Idle boat | 7% | 0% | 6% | ns |
| Other boat | 10% | 8% | 10% | ns |
| Boat | 17% | 24% | 18% | ns |
| Fishing | 5% | 12% | 6% | ns |
| Traffic | 37% | 56% | 40% | ns |
| Train | 2% | 0% | 2% | ns |
| Other noise | 10% | 8% | 10% | ns |
| All anthropophony | 60% | 80% | 63% | ns |
| **Unclassified** |  |  |  |  |
| Bubble-like | 18% | 32% | 20% | ns |
| Gurgle | 21% | 48% | 25% | ** |
| Unknown | 39% | 72% | 43% | ** |
| All unclassified | 46% | 76% | 50% | ** |
| **All Sounds** | 76% | 92% | 79% | ns |
| **Sample size** | 148 | 25 | 173 |  |

Percent of the stations where sound types were observed. P = probability of a significant difference in the frequency of sounds between day and night based on a chisq test (ns = not significant, \* = < 0.05, \*\* = < 0.01, \*\*\* = < 0.001).

Traffic sounds were the most common component of the anthropophony occurring in 37 % of the recordings. Biophony was observed in significantly more recordings at night than during the day (84 % vs 51 %, Table 2). All components of the biophony except insect-like and bird sounds occurred in significantly more night recordings than day recordings. No significant diel differences in occurrence among recordings were observed for the anthropophony.

Other air movement (5 %), FRT (7 %), Other fish (10 %), insect-like (9 %), bird (11 %) and other biological (4 %) sounds overlapped in time with traffic sounds. The only other noises to overlap in time with biological sounds were: airplane (0.1 % and 0.4 %, with other fish and insect-like), other boat (0.1 % and 0.3 %, with other air movement and other fish), and other noise (0.1 % with other fish).

### Diel patterns

The biophony accounted for 66 % and 67 % (mean = 1.7 and 2.7 sounds/min) of the total number of sounds during the day (2.6 sounds/min) and night (4.0 sounds/min), respectively (S7 Fig, Table 3). However, the anthropophony dominated the soundscape in terms of relative percent time accounting for 92 % and 88 % (mean = 15 % and 13 % of the recording time) of total sounds during the day (16.6 % time) and night (14.3 % time), respectively (Fig 3, S7 Fig, Table 3). Unclassified sound accounted for just 9 % of the total sounds by number and less than 5% of the sounds by percent recording time (S7 Fig, Table 3).

**Fig 3.**
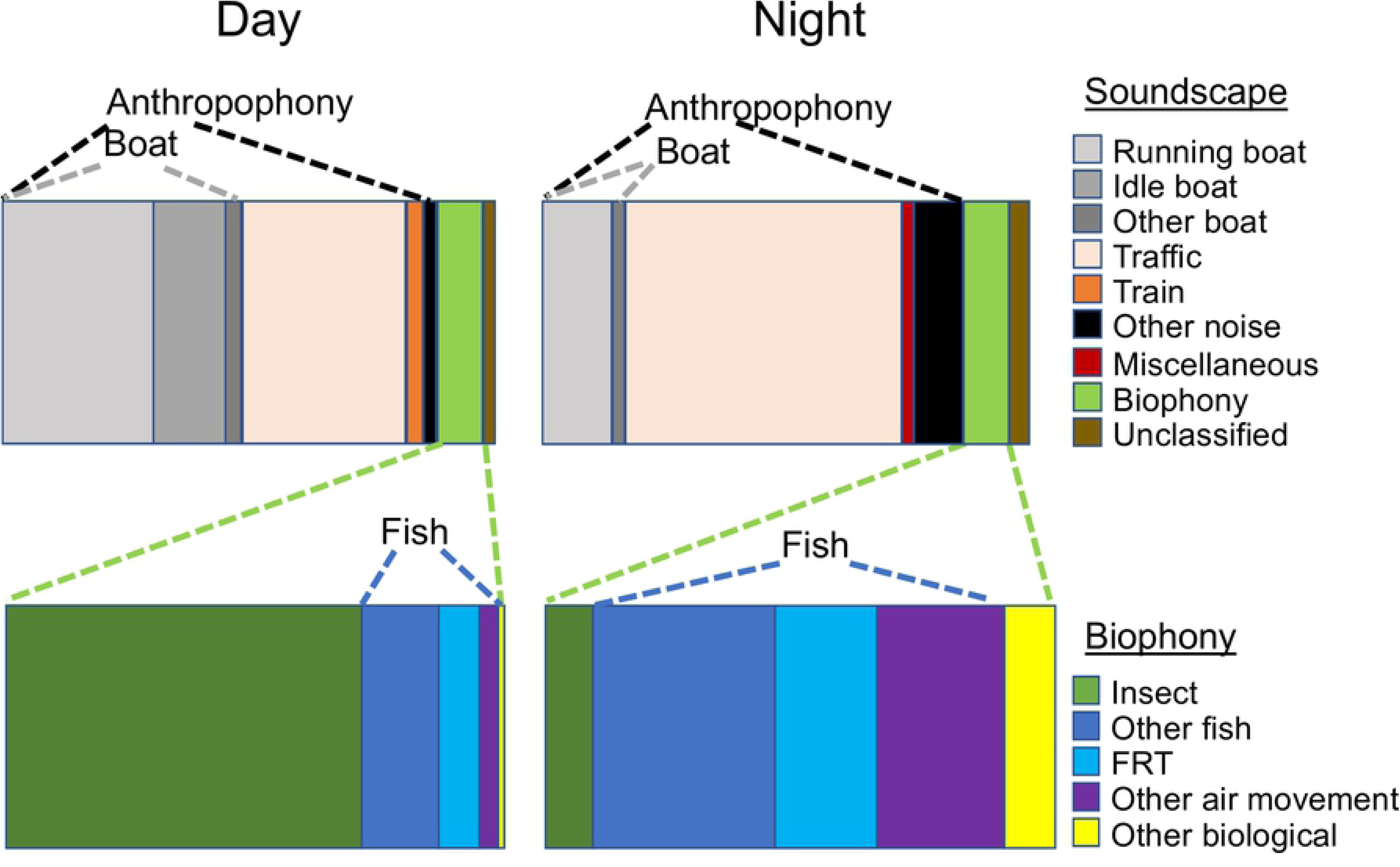
Soundscape composition. Relative contribution of the anthropophony and biophony and their major components to the aquatic soundscape during the day and night based on mean percent time of each sound type (data provided in Table 3). The composition of the biophony is shown in the expanded plots.

**Table 3.** Comparison of sounds between diel periods.

| Sound category | Number per minute |  |  |  |  |  | Percent time |  |  |  |  |  |
| --- | --- | --- | --- | --- | --- | --- | --- | --- | --- | --- | --- | --- |
|  | Day |  |  | Night |  |  | Day |  |  | Night |  |  |
|  | Mean | SE | Max | Mean | SE | Max | Mean | SE | Max | Mean | SE | Max |
| **Biophony** |  |  |  |  |  |  |  |  |  |  |  |  |
| FRT | 0.028 | 0.006 | 0.6 | 0.054 | 0.024 | 0.5 | 0.13 | 0.04 | 4.10 | 0.27 | 0.11 | 1.90 |
| Other air movement | 0.239 | 0.074 | 10.2 | 0.940 | 0.350 | 7.9 | 0.06 | 0.01 | 1.30 | 0.34 | 0.12 | 2.10 |
| Air movement | 0.267 | 0.075 | 10.3 | 0.994 | 0.370 | 8.4 | 0.18 | 0.04 | 4.17 | 0.58 | 0.20 | 4.01 |
| Other fish | 0.585 | 0.201 | 22.3 | 1.316 | 0.790 | 18.9 | 0.24 | 0.08 | 7.50 | 0.48 | 0.22 | 4.90 |
| Fish | 0.852 | 0.221 | 22.3 | 2.310 | 0.913 | 20.2 | 0.44 | 0.10 | 7.50 | 1.09 | 0.33 | 6.30 |
| Insect-like | 0.759 | 0.597 | 88.0 | 0.032 | 0.020 | 0.5 | 1.12 | 0.59 | 76.80 | 0.13 | 0.10 | 2.30 |
| Other biological | 0.076 | 0.019 | 1.5 | 0.198 | 0.064 | 1.1 | 0.02 | 0.01 | 0.40 | 0.14 | 0.09 | 2.10 |
| Surface | 0.029 | 0.010 | 0.9 | 0.111 | 0.042 | 1.0 | 0.04 | 0.01 | 1.38 | 0.14 | 0.08 | 2.04 |
| Birds | 0.015 | 0.004 | 0.4 | 0.017 | 0.008 | 0.1 | 0.04 | 0.02 | 1.76 | 0.07 | 0.05 | 1.08 |
| All biophony | 1.730 | 0.647 | 89.6 | 2.668 | 0.940 | 20.5 | 1.57 | 0.57 | 77.02 | 1.51 | 0.47 | 9.39 |
| **Anthropophony** |  |  |  |  |  |  |  |  |  |  |  |  |
| Airplane | 0.003 | 0.001 | 0.1 | 0.005 | 0.003 | 0.1 | 0.17 | 0.09 | 9.80 | 0.22 | 0.16 | 3.20 |
| Running boat | 0.020 | 0.005 | 0.3 | 0.009 | 0.004 | 0.1 | 5.30 | 1.37 | 92.40 | 2.06 | 0.92 | 16.50 |
| Idle boat | 0.005 | 0.002 | 0.1 | 0.000 | 0.000 | 0.0 | 2.52 | 0.99 | 100.00 | 0.00 | 0.00 | 0.00 |
| Other boat | 0.092 | 0.057 | 8.2 | 0.017 | 0.016 | 0.4 | 0.57 | 0.34 | 46.60 | 0.36 | 0.35 | 8.50 |
| Boat | 0.118 | 0.058 | 8.2 | 0.026 | 0.018 | 0.5 | 8.40 | 1.95 | 100.00 | 2.42 | 1.07 | 16.50 |
| Fishing | 0.009 | 0.004 | 0.4 | 0.045 | 0.025 | 0.5 | 0.04 | 0.02 | 2.10 | 0.11 | 0.07 | 1.70 |
| Traffic | 0.453 | 0.087 | 7.7 | 0.824 | 0.482 | 12.5 | 5.77 | 1.05 | 69.80 | 8.22 | 2.89 | 54.80 |
| Train | 0.008 | 0.005 | 0.6 | 0.000 | 0.000 | 0.0 | 0.56 | 0.41 | 54.90 | 0.00 | 0.00 | 0.00 |
| Other noise | 0.025 | 0.009 | 0.9 | 0.033 | 0.026 | 0.6 | 0.21 | 0.18 | 26.00 | 1.44 | 1.14 | 26.50 |
| All anthropophony | 0.644 | 0.106 | 9.2 | 0.964 | 0.490 | 12.4 | 15.23 | 2.17 | 100.00 | 12.55 | 3.54 | 65.50 |
| **Unclassified** |  |  |  |  |  |  |  |  |  |  |  |  |
| Bubbles | 0.034 | 0.008 | 0.6 | 0.031 | 0.014 | 0.3 | 0.12 | 0.03 | 2.13 | 0.16 | 0.09 | 1.93 |
| Gurgle | 0.045 | 0.009 | 0.7 | 0.199 | 0.104 | 2.5 | 0.10 | 0.03 | 2.42 | 0.34 | 0.18 | 4.10 |
| Unknown | 0.163 | 0.031 | 2.7 | 0.142 | 0.033 | 0.6 | 0.26 | 0.06 | 5.03 | 0.24 | 0.06 | 1.10 |
| All unclassified | 0.242 | 0.035 | 2.8 | 0.371 | 0.127 | 3.0 | 0.47 | 0.08 | 5.03 | 0.71 | 0.22 | 4.92 |
| All sounds | 2.616 | 0.660 | 89.6 | 4.002 | 1.166 | 25.4 | 16.55 | 2.12 | 100.00 | 14.26 | 3.58 | 66.02 |
| **Sample size** | 148 |  |  | 25 |  |  | 141 |  |  | 24 |  |  |

Mean number per minute and percent of recording time of selected sound categories for day and night sampling. SE = standard error of the mean, Max = maximum value.

Biological sounds tended to be more diverse at night averaging 7.9 types per recording (standard error = 1.7, maximum = 32) compared to 2.5 types per recording (standard error = 0.3, maximum = 16) types during the day. Although the relative contribution of biophony and anthropophony by both number and percent recording time were similar between day and night, the composition of the sounds changed (Fig 3, S7 Fig, Table 3). Insect sounds dominated the biophony by day, while total fish sounds dominated by night. Air movement fish sounds and other fish sounds contributed about equally to total fish sounds, though other fish sounds were more numerous during the day, while the longer duration air movement sounds accounted for more of the recording time at night. Traffic sound was the most numerous component of the anthropophony during both day and night, but the long-duration boat sounds contributed more to the recording time during the day. Traffic sound accounted for more than half of the recorded sound during the night based on mean percent of recording time.

### Habitat patterns

There were no significant differences in the percent of recordings among habitat categories within the biophony during the day (S3 Table). At night, insect sounds occurred more frequently in the brook/creek habitat (however, the brook/creek sample size was low), while other biological and bird sounds were absent from the pond/lake habitat. Traffic sounds were the most widespread noise and were significantly more frequently occurring in brook/creek habitat during the day. In contrast, boat sounds were absent from brook/creek (S3 Table). No significant differences in the frequency of occurrence of anthropophonic sounds were observed among the habitat categories at night (S3 Table).

Insect and other fish sounds dominated the biophony by percent time in all habitats during the day, but air movement sounds dominated pond/lake and river habitats at night (S8 Fig, S4 Table). Other fish dominated brook/creek habitat at night but the sample size was low (n = 2). Traffic sounds dominated soundscape percent time during both day and night in brook/creek habitat, while boat sounds dominated other habitats (S8 Fig, S4 Table).

### River gradient pattern

There were no consistent trends among rivers in biological or noise sounds in main-stem river habitats moving along the river gradient from headwaters to mouth, although the highest elevation locations tended to have little or no biological sounds (S1 Database S1 online). When day data from all rivers were pooled after grouping locations into non-tidal (N = 46) and tidal regions (N = 20), all boat noise categories were significantly more frequent in tidal regions (S5 Table). Similar trends were observed for sound rate and percent time (S9 Fig, S5 Table). Boat noise averaged 31 % of the recording time in tidal regions, but only 2 % in non-tidal areas. Similarly, boat sounds (all types combined) averaged 0.02 sounds/min and 0.21 sounds/min in non-tidal and tidal regions, respectively (P < 0.0001). Average soundscape percent time of traffic sounds and all anthropophonic sounds combined were also significantly different between river regions (traffic = 8.3 % and 0.8 %, noise = 12.1 % and 33.2 %, for non-tidal and tidal regions, respectively, both P < 0.05). Surface sounds were significantly more frequent, numerous, and occupied more time in tidal regions than non-tidal. No other biophonic sounds were significantly different among regions (S9 Fig, S5 Table).

### Ambient sound level

Overall received RMS values ranged from 90 to 133 dB re 1 μPA. There was no significant difference in received RMS level among habitat types during the day with averages (and standard errors) of 99.4 (2.2), 98.7 (1.2) and 101.1 (1.4) dB re 1 μPA, for brook/creek, pond/lake and river habitats, respectively (based on a one-way ANOVA on log transformed data; N = 77). However, average spectra of the ambient sounds suggest differences in the frequency structure among habitat types (S6 Fig). Brook/creek habitats tend to have the highest levels and pond/lake habitats the lowest levels at frequencies below 500 Hz, while river habitats have the highest levels at all higher frequency bands.

Background ambient sound levels (RMS category) had a strong influence on biological sound occurrence (Fig 4, S7 and S8 Tables). Air movement sounds significantly declined from a high of 72 % of recordings to a low of 6 % of recordings from the lowest to highest ambient sound level categories. Similar, but non-significant, trends in occurrence were observed for FRT and other fish sounds (Fig 4). Rate and percent time of biological sounds followed similar trends with significant declines in air movement, fish and total biophony with increasing RMS level (S7 Table). Biodiversity also declined significantly from 4.2 to 0.7 sounds/recording with increasing ambient sound level (S7 Table).

**Fig 4.**
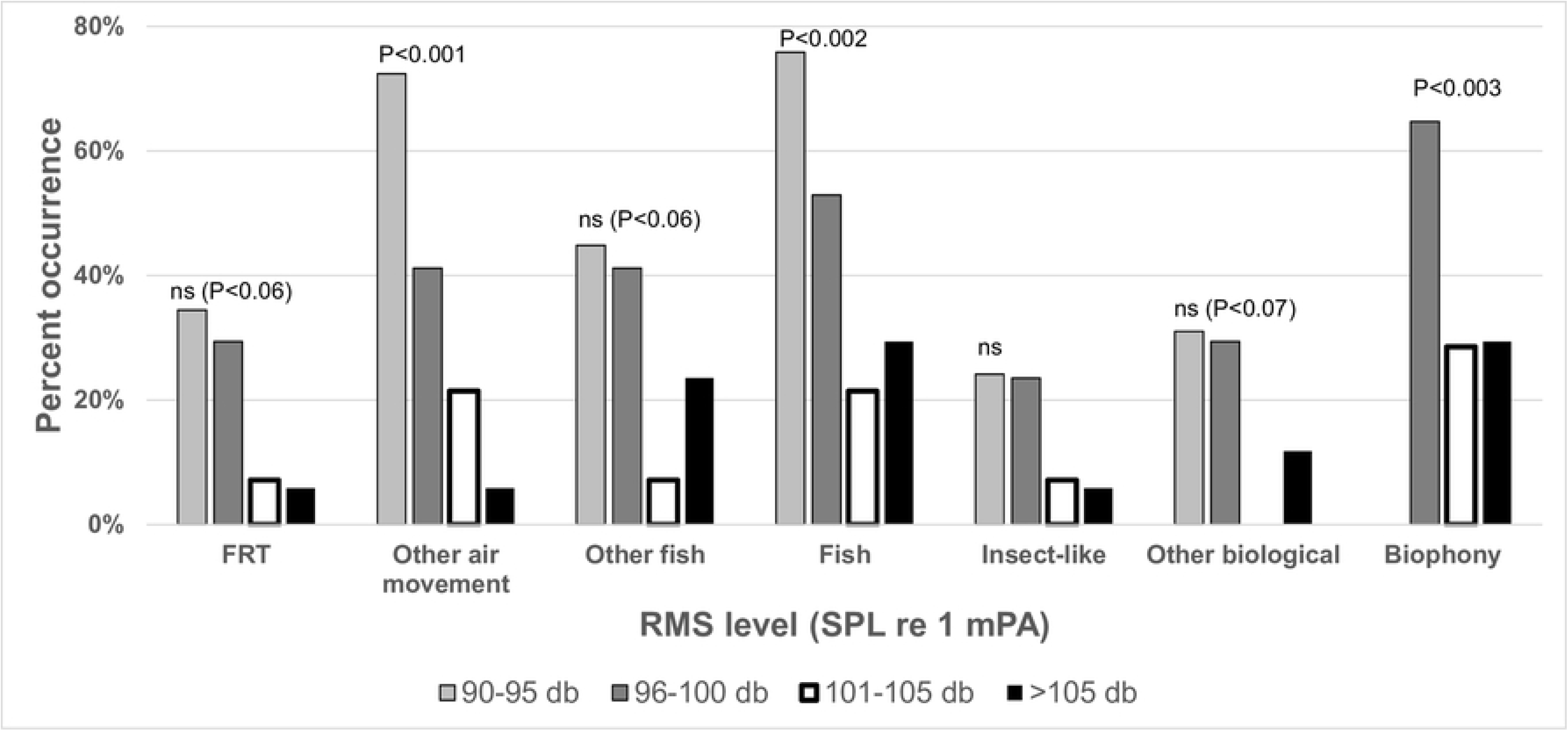
Ambient sound level effect. Comparison of the frequency of occurrence of selected biophony components among locations grouped into four ambient sound levels based on received root mean square (RMS) measurements (N = 77, see S6 Table for details). Significant differences in the observed group frequency from the expected group frequency are indicated (P ≤ 0.05, ns = not significance).

A comparison of the average power spectra of major biophony categories with major anthropophony categories indicates that other fish sounds are above average anthropogenic sounds, except for running and other boat sounds (S6 Fig). There is also significant overlap with the spectra of traffic sounds. Air movement and insect sounds produce low amplitude sounds largely below anthropogenic sound levels and ambient spectra averaged across all recordings (S6 Fig), but which nevertheless exhibit significant energy above ambient noise from the locations they occurred at, except for insect sounds which were often close to ambient levels (Fig 5).

**Fig 5.**
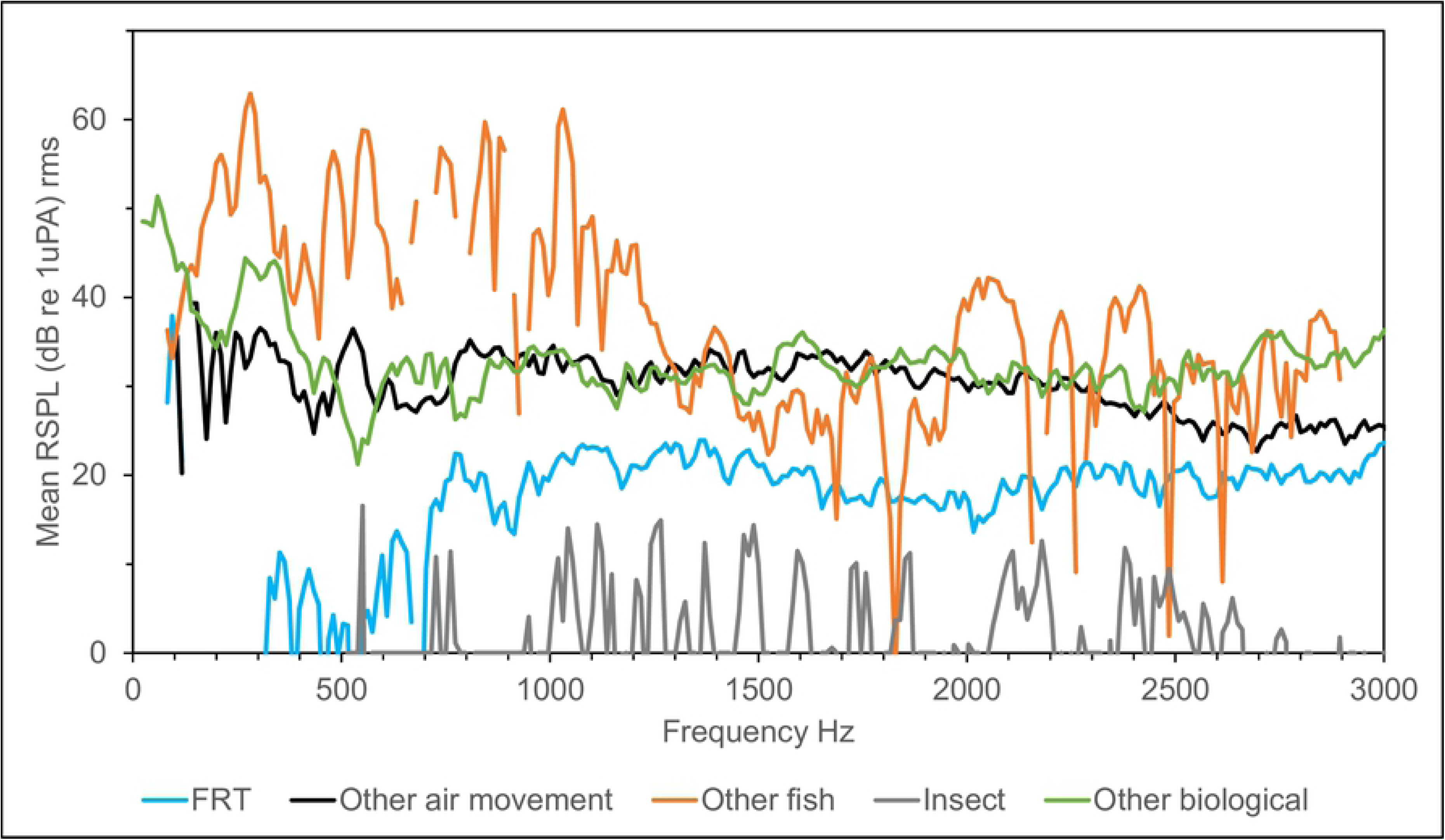
Selected biophony spectra above local ambient sound. Mean power spectra above ambient of selected biophony categories, calculated as the received level difference between individual sounds and the ambient spectra from the same recording and then averaging among all subsampled sounds within each biophonic category (Hanning, FFT 4096, 50% overlap, frequency resolution 11.7, PSD normalized).

## Discussion

The three most significant findings in this study are: 1) it builds upon our previous study of the Hudson River [12] to demonstrate that freshwater habitats in the New England region of North America contain a diverse array of as yet unidentified biological sounds; 2) together with our companion paper [10], it demonstrates for the first time that air movement sounds constitute an important component of the freshwater soundscape; and 3) anthropogenic noises now dominate the soundscape and their overlap in time and frequency with the biophony suggests a high potential for negative impacts. The impact of anthropogenic noise on the natural freshwater soundscape is illustrated in Figs 3 and 4, which indicate that the anthropophony accounts for about 90 % of the soundscape, and that biophony declines (or is masked from detection) with increasing ambient sound levels (see also S10 Fig).

Although several recent studies have examined the diversity of ambient noise and potential impacts on specific fishes [3-7, 15-16 and citations therein), none have quantified the relative composition of both biophony and anthropophony components of the soundscape, but rather have focused on quantification of noise levels [except 7]. We suggest that quantification of the soundscapes in terms of the percent time of component sounds is a useful way to compare habitats. Although percent time does not provide information on sound level, it taken together with information on temporal and frequency overlap of biophony with anthropophony does provide useful information on the exposure to noise and likelihood of negative impacts. Consider that any level of anthropophonic noise in these habitats alters the soundscape from that in which aquatic organisms have evolved in, and if the frequency of the noise overlaps with either the hearing range or sound production bandwidth of specific organisms, then there is a potential for negative impacts of noise on those organisms.

Our observations of the widespread occurrence of air movement sounds in many habitats across a large geographic area, together with their large contribution to the biological soundscape based on sound rate and sound percent time, suggest for the first time that air movement sounds are an important phenomenon in freshwater habitats. We emphasize that even if air movement sounds are largely incidental, if sounds are species specific [10], they can be used by scientists and resource managers as an aid in documenting the spatial and temporal distribution of fishes and their associated soniferous behavior [17]. In addition, we point out that even incidental sounds can expose an organism to predation by organisms that can hear that sound, and hence would be subject to natural selection pressures.

Despite the lack of significant habitat differences in ambient sound levels, sound level appeared to have a strong negative influence on biological diversity and soundscape composition (Fig 3, S6 and S7 Tables), suggesting possible masking, suppression of sound production, or avoidance of locations with high ambient noise levels. A negative effect of ambient noise level on biological sound production or detection supports previous work on potential impacts of noise levels on fishes [e.g. 3-7, 15-16], however we caution that the results may have been biased by confounding effects since no biological sounds were negatively correlated with any noise rate or percent time variable. The lack of such correlations may simply be due to the high diversity of sounds observed while sampling over a wide variety of habitats and geographies with many different faunal assemblages. The overlapping frequency structure of noises, especially with the other fish category (Fig 2, S6 Fig, Table 1) also suggests the potential for masking (an example of which can be seen in S1 Fig and the corresponding S1 Audio). In contrast, the lack of overlap between peak frequencies of biological sounds and peak ambient frequency (Fig 2, S6 Fig) lends further support to previous studies suggesting that fish take advantage of acoustic niches [18]; however past studies have described low frequency acoustic windows in shallow fast-flowing streams [18]. Our data suggest that a high-frequency “quiet noise window” may also occur in other freshwater habitats, and that freshwater organisms may have evolved to exploit different acoustic windows in different habitats. It is also possible that multiple acoustic windows may occur in some habitats which can be exploited by different organisms.

Running boats generate long lasting noise starting with low-amplitude and low-frequency noise while in the distance and progressing to high amplitude noise that saturates the recordings when within a few hundred meters (depending on the boat type and speed), before gradually fading as the boat moves off in the distance again (an example of the sound of an approaching boat which anchors within 100 m of the recording site is provided in S3 Fig and corresponding S3 Audio). Over the years we have frequently observed boat noise percent time at 100 % in some recordings, particularly in estuarine habitats (Rountree et al. *Pers. Observ*.). Locations near important navigation routes, marinas, or boat landings experience chronic boat noise exposure [e.g., 19]. In some cases, boat noise is so prevalent in the hours around sunset (when boaters are returning to shore) that recording of biological sound is nearly impossible. We point out that our sampling was conducted during the early season when boating activity was sparse and many boat ramps had not yet opened for the season. We would expect, therefore, that boat noise would be significantly more prevalent during the summer months than that observed in our study. Boat noise is undoubtedly an important chronic noise source in navigable waters where it dominates the soundscape and unquestionably masks many biological sounds (S4 and S6 Figs). Boat noise is particularly problematic in enclosed water bodies such as small lakes, and in linear rivers as the sound travels great distances. We have often detected motor boat sounds before sighting the vessel in the distance. On the other hand, serpentine waterways may be less impacted because sounds of an approaching vessel are blocked by land until the vessel moves around the bend and into the line of sight.

Our observations suggest that boats running idle while docked, anchored, or drifting are a major component of freshwater soundscapes (Fig 3), and have the potential to mask some biophony such as insect sounds, but we are not aware of studies that examine its potential impact. The tendency for boaters, and especially ferries and other large vessels, to idle for long periods, and the lower frequency structure of idling boats (Fig 2, S7 Fig) suggests that boat idling may be an important chronic noise source in navigable waters.

Traffic noise exhibits similar though much less extreme wax-and-wane patterns as can clearly be seen in S1 Fig and heard in the corresponding S1 Audio. Our observations support those of Holt and Johnston [3] that traffic sounds can have an important impact on some freshwater habitats. Although not as extreme as boat noise, traffic sounds are far more ubiquitous in freshwater habitats and can be chronic in urban areas and during rush hours. In fact, the increase in traffic contribution to the soundscape at night was due in part to our sampling around sunset when traffic tends to be heavy. The relatively high temporal overlap between biological sounds and traffic sounds suggests a high likelihood for impacts, especially for other fish sounds which had both high temporal (10 %) and acoustic frequency overlap (Fig 2, S6 Fig). Although we did not examine propagation distances of traffic sounds, traffic crossing bridges was detected in locations as far as 270 m from the nearest bridge (S1 Data Base), agreeing well with previous studies [3, 5]. Differences in the relative contribution of traffic and boat noises to the soundscape among habitats and river regions demonstrate variations in potential noise impacts among habitats and river zones (S9 Fig, S5 Table). It should be recognized that traffic sounds were likely over represented in our sampling due to the frequent necessity of accessing the water near bridges. However, as has previously been pointed out [3], in many areas bridge crossings are common and smaller streams and rivers may be crossed many times over short stretches in urban and suburban locations.

The inconsistent trends in observations moving along the river main stem locations likely results from the fact that river systems form a coenocline where gradients in abiotic and biotic conditions regulate community structure and habitat function [20]. Thus, comparison among river systems of different lengths and elevation gradients requires a gradient approach. It is interesting that while the biotic community changes considerably from high elevation reaches to estuarine reaches, the changes in the biophony type contribution to the soundscape are minimal, suggesting that although soniferous species may change, the broad sound categories are more consistent. Sampling along the river coenocline at a higher spatial resolution would likely reveal more subtle shifts in the biophony, due to species assemblage changes. A gradient in impacts from different types of anthropogenic noise is expected as we observed a striking transition from remote wilderness to increasingly developed urban areas while traveling from the river headwaters to the sea. Some of this transition was captured in the comparison between non-tidal and tidal reaches of the rivers where there was a shift between dominance of the soundscape by traffic noise in non-tidal reaches to a dominance by boat noise in tidal reaches (S9 Fig, S5 Table), highlighting different potential for impacts in different habitats.

Why has the freshwater soundscape been neglected by scientists, resource managers, conservationists and environmentalists for so long? We believe that in large part it is due to a general lack of appreciation of the importance of inland fisheries on regional, national and international scales which can lead to a lack of scientific inquiry [21]. A lack of appreciation of the importance of the soundscape is exacerbated by a pervasive lack of awareness that fish and other aquatic organism produce sounds that can be monitored remotely and how that can be a valuable tool for scientists and resource managers [17]. Similarly, fisheries scientists, resources managers and conservationists have been slow to understand the importance of the soundscape to non-vocal organisms [22–23] (but see [24] for ecosystem effects of noise on non-vocal marine invertebrates). Although there has been a dramatic increase in awareness and attention to soundscapes in marine systems in the last decade, and in particular the effects of noise [25], attention has only recently begun to trickle over to freshwater systems. Furthermore, the lack of familiarity with biological sounds in freshwater habitats [8] contributes to their being overlooked.

## Acknowledgments

Joe Olson provided technical assistance. Megan O’Connor provided the map used in Fig 1.

## Supplemental materials

**S1 Fig. Representative sampling sites.** A) Brook wp 42, B) Creek wp 60, C) Pond wp 68, D) Lake wp 83, E) Tributary wp 117, F) River wp 101. The waypoint (wp) number can be used to look up the location details in the S1 Data set.

**S2 Fig. Example of traffic sound**. Relative amplitude (top) and spectrogram of traffic and fish sounds recorded at night on 14 May 2008 in Sucker Creek, Griffin, New Hampshire (N43° 00.257’ W71° 20.933’). As a car passes over a nearby bridge (Traffic), catfish sounds (Other Fish) are partially masked. An unmasked catfish sound occurs later in the clip and indicates a true peak frequency well within the traffic noise. Examples of a fish splashing at the surface (Surface), and subsequent air movement (Other Air movement) and FRT sounds can also be seen relative to the traffic sound. An amplified audio file corresponding to the figure can be heard in S1 Audio online. Spectrogram parameters: unfiltered, 1,024-point Hann windowed FFTs with 50% overlap.

**S3 Fig. Example of train noise**. Recorded on 3 May 2008 in a tributary of the Connecticut River in Deep River, Connecticut (N41° 22.978’ W72° 25.566’). Top: photograph of the train passing by at the time of the recording. Bottom: relative amplitude waveform and spectrogram of the train sound which can be heard in the corresponding S2 Audio online. Spectrogram parameters: unfiltered, 1,024-point Hann windowed FFTs with 50% overlap.

**S4 Fig. Example of running boat noise.** The sound of an outboard motor boat as it approaches from the distance and stops to anchor nearby, recorded on 3 May 2008 in the mainstem of the Connecticut River in Old Saybrook, Connecticut (N41° 19.143’ W72° 21.028’). Top: Photograph of the boat as it passes, Bottom: relative amplitude waveform and spectrogram of the noise generated by the passing boat, which can be heard in the corresponding S3 Audio online. Spectrogram parameters: unfiltered, 1,024-point Hann windowed FFTs with 50% overlap.

**S5 Fig. Example of other boat noise.** Sound produced by the power trim of a nearby outboard boat, recorded on 1 May 2008 in the mainstem of the Connecticut River in Northampton, Massachusetts (N42° 20.114’ W72° 37.211’). The corresponding sound can be heard in the S4 Audio online. Spectrogram parameters: unfiltered, 1,024-point Hann windowed FFTs with 50% overlap.

**S6 Fig. Average power spectra of major sound categories.** Power spectra averaged over a subsample of sounds from each major sound category (samples sizes are shown in S2 Table). A) selected biophonic sounds, B and C) selected anthropophonic sounds, D) ambient sounds from each habitat category. Spectrogram parameters: Hanning, FFT 4096, 50% overlap, frequency resolution 11.7, PSD normalized.

**S7 Fig. Comparison between day and night soundscapes.** Major components of the soundscape: A and C) during the day and night, respectively, based on mean number, B and D) during day and night, respectively, based on mean percent time. The relative contribution of the biophony compared to the anthropophony is shown in the main pie, while the composition of the biophony slice is shown in the expanded pie. The size of the wedge represents the relative proportion of the sound out of all sounds based on data found in Table 2 of mean number per minute (A and C) and mean percent time (B and D).

**S8 Fig. Comparison of habitat soundscapes.** Comparison of soundscapes among habitat categories and diel period based on mean percent time. Day: A) creek/brook habitat during the day (N = 21), B) pond/lake habitat (N = 41), C) river habitat (N = 79). Night: D) creek/brook habitat (N = 2), E) pond/lake habitat (N = 7), F) river habitat (N = 15). See Supplementary Table S4 for summary statistics.

**S9 Fig. Comparison of soundscapes between tidal and non-tidal river regions.** Relative contribution of anthropophony and biophony to the aquatic soundscape of non-tidal (top) and tidal (bottom) main-stem river regions during the day based on mean number of sounds per minute (A and B) mean percent time (C and D) of each sound type. Data and statistics are provided in S5 Table, while the size of the wedge represents the relative proportion of the sound out of all sounds.

**S10 Fig. Average day-time soundscape composition.** Venn diagram illustrating the relative composition of the anthropophony and biophony and their major constituents to the day-time soundscape. The diameter of each circle is proportional to the mean percent of recording time for the indicated sound category. Circles within large circles represent subcomponents of the larger category. For example, the fish category contains two nearly equal subcomponents (other fish and air movement sound).

S1 Table. Sampling effort for all locations.

S2 Table. Subsample sizes used for spectral analysis by sound type.

**S3 Table. Percent occurrence of sounds at locations by habitat category.** (P = significance level for a Chisq test for differences among habitats within a diel period, * = < 0.05, ** = < 0.01, *** = <0.001, n/a = no test; N = 148 for day and 25 for night).

**S4 Table. Mean percent time by habitat category**. Comparison of the mean percent time (total duration of sounds in the category divided by recording duration) of sound types among habitat types. SE = standard error of the mean. P = probability of a significant difference among habitats based on a one-way ANOVA on transformed variables, performed separately by diel period. (* = < 0.05, ** = < 0.01, *** = <0.001, n/a = no test).

**S5 Table. Comparison of summary statistics between tidal and non-tidal river regions.** Mean number per minute, percent of recording time, and percent occurrence of sound categories for day sampling within tidal and non-tidal river regions (SE = standard error of the mean, P = significance level from a one-way analysis of variance, Chisq P = significance level from a chisq test of expected frequencies, * ≤ 0.05, ** ≤ 0.01, *** ≤ 0.001).

**S6 Table. Frequency of occurrence of biophony among ambient sound level categories.** Comparison of the percent frequency of occurrence among day-time recordings grouped into received ambient sound level categories (RMS = root mean square). (P = significance level from a chisq test of difference among RMS categories from the expected frequency. ns = not significant, * ≤ 0.5, ** ≤ 0.01, *** ≤ 0.001).

**S7 Table. Comparison of the mean number and mean percent time among daytime ambient sound categories.** (P = results of a one-way ANOVA on differences among RMS levels of transformed variables. ns = not significant, * ≤ 0.5, ** ≤ 0.01, *** ≤ 0.001).

**S1 Audio. Traffic sound with biophony.** Sound file corresponding to S2 Fig containing the sound of traffic noise and examples of other fish, other air movement and FRT sounds recorded at night on 14 May 2008 in Sucker Creek, Griffin, New Hampshire (N43° 00.257’ W71° 20.933’). The sound has been amplified for optimal listening online.

**S2 Audio. Train sound.** Example of train noise corresponding to S3 Fig and recorded on 3 May 2008 in a tributary of the Connecticut River in Deep River, Connecticut (N41° 22.978’ W72° 25.566’). Note that several very faint other air movement sounds (chirps) can be heard in the recording but are high frequency and not shown in the S3 Fig. The sound has been amplified for optimal listening online.

**S3 Audio. Running boat sound.** Example of the sound of an outboard motor boat, corresponding to S4 Fig, as it approaches from the distance and stops to anchor nearby. Recorded on 3 May 2008 in the mainstem of the Connecticut River in Old Saybrook, Connecticut (N41° 19.143’ W72° 21.028’). The sound has been amplified for optimal listening online.

**S4 Audio. Other boat noise.** Example of noise produced by the power trim of a nearby outboard boat, corresponding to S5 Fig. Recorded on 1 May 2008 in the mainstem of the Connecticut River in Northampton, Massachusetts (N42° 20.114’ W72° 37.211’). The sound has been amplified for optimal listening online.

**S1 Data set. Meta-data of sounds observed in all recordings.** Recording location meta-data with sound rates and percent time by sound category. The data set is in Excel format with two worksheets: 1) data, 2) field definitions.

## References

1. Abbott CC. Traces of a voice in fishes. The American Naturalist. 1877;11: 147–156.

2. Stober QJ. Underwater noise spectra, fish sounds and response to low frequencies of Cutthroat trout (*Salmo clarki*) with reference to orientation and homing in Yellowstone Lake. Trans. Am. Fish. Soc. 1969;98(4): 652–663.

3. Tonolla D, Acuna V, Lorang MS, Heutschi K, Tockner K. A field-based investigation to examine underwater soundscapes of five common river habitats. Hydrological Processes 2010;24(22); 3146–3156.

4. Holt DE, Johnston CE. Traffic noise masks acoustic signals of freshwater stream fish. Biological Conservation 2015;187: 27–33.

5. Bolgan M, Chorazyczewska E, Winfield IJ, Codarin A, O’Brien J, Gammell M. First observations of anthropogenic underwater noise in a large multi-use lake. Journal of Limnology. 2016 Jun 14;75(3): 644–651.

6. Martin SB, Popper AN. Short- and long-term monitoring of underwater sound levels in the Hudson River (New York, USA). J. Acoust. Soc. Am. 2016;139(4): 1886–1897.

7. Bolgan M, O’Brien J, Chorazyczewska E, Winfield IJ, McCullough P, Gammell M. The soundscape of Arctic Charr spawning grounds in lotic and lentic environments: can passive acoustic monitoring be used to detect spawning activities? Bioacoustics. 2018 Jan 2;27(1): 57–85. http://dx.doi.org/10.1080/09524622.2017.1286262

8. Rountree RA, Bolgan M, Juanes F. How Can We Understand Freshwater Soundscapes Without Fish Sound Descriptions? Fisheries. 2019 Mar;44(3):137–43. https://doi.org/10.1002/fsh.10190

9. Rountree RA, Juanes F. Potential of passive acoustic recording for monitoring invasive species: freshwater drum invasion of the Hudson River via the New York canal system. Biological Invasions. 2017 Jul 1;19(7):2075–88.

10. Rountree RA, Juanes F, Bolgan M. Air movement sound production by alewife, white sucker, and four salmonid fishes suggests the phenomenon is widespread among freshwater fishes. PloS one. 2018 Sep 20;13(9):e0204247. doi: 10.1371/journal.pone.0204247.

11. Johnson NS, Higgs D, Binder TR, Marsden JE, Buchinger T, Brege L, Bruning T, Farha S, Krueger CC. Evidence of sound production by spawning lake trout (*Salvelinus namaycush*) in lakes Huron and Champlain. Canadian Journal of Fisheries and Aquatic Sciences. 2017 May 5;75(3):429–38. doi: 10.1139/cjfas-2016-0511.

12. Anderson KA, Rountree RA, Juanes F. Soniferous fishes in the Hudson River. Trans. Am. Fisher. Soc. 2008;137(2): 616–626.

13. Hawkins AD, Popper AN. Assessing the impacts of underwater sounds on fishes and other forms of marine life. Acoust. Today 2014;10(2): 30–41.

14. Charif RA, Waack AM, Strickman LM. Raven Pro 1.4 user’s manual. Cornell Lab of Ornithology, Ithaca, NY. 2010 May 22;25506974.

15. Wysocki LE, Amoser S, Ladich F. Diversity in ambient noise in European freshwater habitats: Noise levels, spectral profiles, and impact on fishes. The Journal of the Acoustical Society of America. 2007 May;121(5):2559–66.

16. Amoser S, Ladich F. Year-round variability of ambient noise in temperate freshwater habitats and its implications for fishes. Aquatic Sciences. 2010 Jun 1;72(3):371–8.

17. Rountree RA, Gilmore RG, Goudey CA, Hawkins AD, Luczkovich JJ, Mann DA. Listening to fish: applications of passive acoustics to fisheries science. Fisheries. 2006 Sep 1;31(9):433–46.

18. Lugli M. Sounds of shallow water fishes pitch within the quiet window of the habitat ambient noise. Journal of Comparative Physiology A. 2010 Jun 1;196(6):439–51.

19. Correa JM, Sempere JT, Juanes F, Rountree R, Ruíz JF, Ramis J. Recreational boat traffic effects on fish assemblages: First evidence of detrimental consequences at regulated mooring zones in sensitive marine areas detected by passive acoustics. Ocean & coastal management. 2019 Feb 1;168:22–34. DOI: 10.1016/j.ocecoaman.2018.10.027.

20. Rountree RA, Able KW. Spatial and temporal habitat use patterns for salt marsh nekton: implications for ecological functions. Aquatic Ecology. 2007 Mar 1;41(1):25–45.

21. Cooke SJ, Allison EH, Beard TD, Arlinghaus R, Arthington AH, Bartley DM, Cowx IG, Fuentevilla C, Leonard NJ, Lorenzen K, Lynch AJ. On the sustainability of inland fisheries: finding a future for the forgotten. Ambio. 2016 Nov 1;45(7):753–64.

22. Cotter AJ. The ‘soundscape’of the sea, underwater navigation, and why we should be listening more. Advances in Fisheries Science. 2008 Apr 17;50:451–71.

23. Fay R. Soundscapes and the sense of hearing of fishes. Integrative Zoology. 2009 Mar;4(1):26–32.

24. Solan M, Hauton C, Godbold JA, Wood CL, Leighton TG, White P. Anthropogenic sources of underwater sound can modify how sediment-dwelling invertebrates mediate ecosystem properties. Scientific reports. 2016 Feb 5;6:20540.

25. Cox K, Brennan LP, Gerwing TG, Dudas SE, Juanes F. Sound the alarm: A meta-analysis on the effect of aquatic noise on fish behavior and physiology. Global Change Biology. 2018 Jul;24(7):3105–16. DOI: 10.1111/gcb.14106.

